# Causal Contributions of Left Inferior and Medial Frontal Cortex to Semantic and Executive Control

**DOI:** 10.1101/2025.04.16.649200

**Authors:** Sandra Martin, Matteo Ferrante, Andrea Bruera, Gesa Hartwigsen

## Abstract

Semantic control guides the targeted and context-based retrieval from semantic memory. The overlap with and dissociation from domain-general executive control in the frontal lobe remains contentious. Here, we used transcranial magnetic stimulation (TMS) to probe the functional relevance of the left inferior frontal gyrus (IFG) and pre-supplementary motor area (pre-SMA) for semantic and executive control. Across four sessions, 24 participants received 1 Hz repetitive TMS to each region individually, dual-site TMS targeting both regions sequentially (IFG followed by pre-SMA), and sham TMS. Participants then completed semantic fluency, figural fluency, and picture-naming tasks. Stimulation of either region broadly disrupted both semantic and figural fluency, suggesting shared functionality. However, electric field modeling of the induced stimulation strength revealed distinct specializations: The left IFG was primarily associated with semantic control, as evidenced by verbal fluency deficits, while the pre-SMA played a domain-general role in executive functions, affecting non-verbal fluency and cognitive flexibility (e.g., clustering and switching during semantic fluency). Notably, only dual-site TMS impaired accuracy in figural fluency, providing unique evidence for successful compensation of executive functions through either the left IFG or pre-SMA following single-site perturbation. These findings underscore the multidimensionality of cognitive control and suggest a flexible task-dependent contribution of the IFG to control processes, either as semantic-specific or general executive resource. Furthermore, they highlight the tightly interconnected network of executive control subserved by the left IFG and pre-SMA, advancing our understanding of the neural basis of semantic and executive functions.

## Introduction

Semantic cognition—the ability to flexibly access and use knowledge—relies on two key components: semantic memory, which stores concepts and facts, and semantic control, which guides the targeted retrieval based on context. Neuroimaging and lesion studies reveal distinct yet interacting brain networks supporting these processes (Jefferies, 2013; Jefferies et al., 2008). Semantic memory is represented by a distributed network of modality-specific areas, reflecting all the different information relevant to semantic concepts, and a transmodal hub in the anterior temporal lobe, which bridges information across modalities (Chiou et al., 2018; Lambon Ralph et al., 2017). This network is organised bilaterally, reflecting the broad distribution of conceptual knowledge across the cortex. In contrast, semantic control engages a primarily left-lateralized network involving frontal and temporal regions. The left posterior middle temporal gyrus and the left inferior frontal gyrus (IFG) have emerged as key regions of the semantic control network (Jackson, 2021; Jefferies et al., 2008; Noonan et al., 2013). Current debates center on the involvement of domain-specific versus domain-general control networks in semantic processing (Chiou et al., 2023; Diveica et al., 2023; Jung & Lambon Ralph, 2023). Here, we address this question by probing the functional relevance of two key regions of semantic and domain-general executive control in verbal and non-verbal fluency tasks.

The involvement of domain-general executive control in semantic processing remains contentious. A recent meta-analysis on the neural correlates of semantic control unveiled a primarily specialized network for semantic control, while also identifying overlapping regions with the domain-general multiple-demand network (MDN), specifically in the pre-supplementary motor area (pre-SMA) and the left IFG (Jackson, 2021). Functional neuroimaging studies have revealed a gradient within these areas, indicating a functional subdivision between semantic-specific and domain-general control regions (Chiou et al., 2023; Diveica et al., 2023; Fedorenko et al., 2012). However, studies that explicitly modulated the cognitive and semantic demand of a task showed increased MDN activity under high task loads, suggesting at least a supportive role of the MDN in semantic control (Gao et al., 2021; Martin et al., 2022; Nieberlein et al., 2024).

Fluency tasks offer a unique opportunity to explore the interplay between domain-specific and domain-general executive control as they rely on executive functions such as processing speed, updating, and inhibition (Aita et al., 2019; J. Amunts et al., 2020), while also engaging distinct domain-specific control processes. For example, semantic fluency, a common neuropsychological test, requires the goal-directed navigation of semantic memory for controlled lexical retrieval (Shao, 2014; Whiteside et al., 2016). In contrast, figural fluency, which requires participants to create original line designs, engages visuospatial and visuomotor skills, and is considered an important test of non-verbal fluency (Foster et al., 2005).

In this study, we used transcranial magnetic stimulation (TMS) to investigate the causal roles of the pre-SMA and left anterior IFG in semantic-specific and domain-general executive control. We applied offline repetitive TMS (rTMS) to these regions individually and in combination using a dual-site stimulation approach, followed by semantic and figural fluency tasks (Fig. 1). Comparing the effects of single-site and dual-site TMS allowed us to investigate the potential joint contribution of both areas to fluency performance and their capacity to compensate for perturbation of the other area (Hartwigsen et al., 2010). We hypothesized that IFG perturbation would selectively impair semantic fluency, while pre-SMA disruption would affect both tasks. Combined stimulation was expected to have a stronger effect on semantic fluency if these regions jointly contribute to the process. Using electrical field (e-field) modeling, we explored the relationship between stimulation strength and behavioral changes, and associated the TMS effect with subregions in the stimulated cortical areas. Additionally, we employed a novel machine learning-based clustering and switching analysis to examine TMS effects on category switching during verbal fluency, a process highly sensitive to the detection of neurodegenerative diseases (Troyer et al., 1998). Given the executive nature of switching performance, we anticipated a relatively stronger impact of pre-SMA and dual-site TMS compared to IFG stimulation alone.

**Figure 1.**
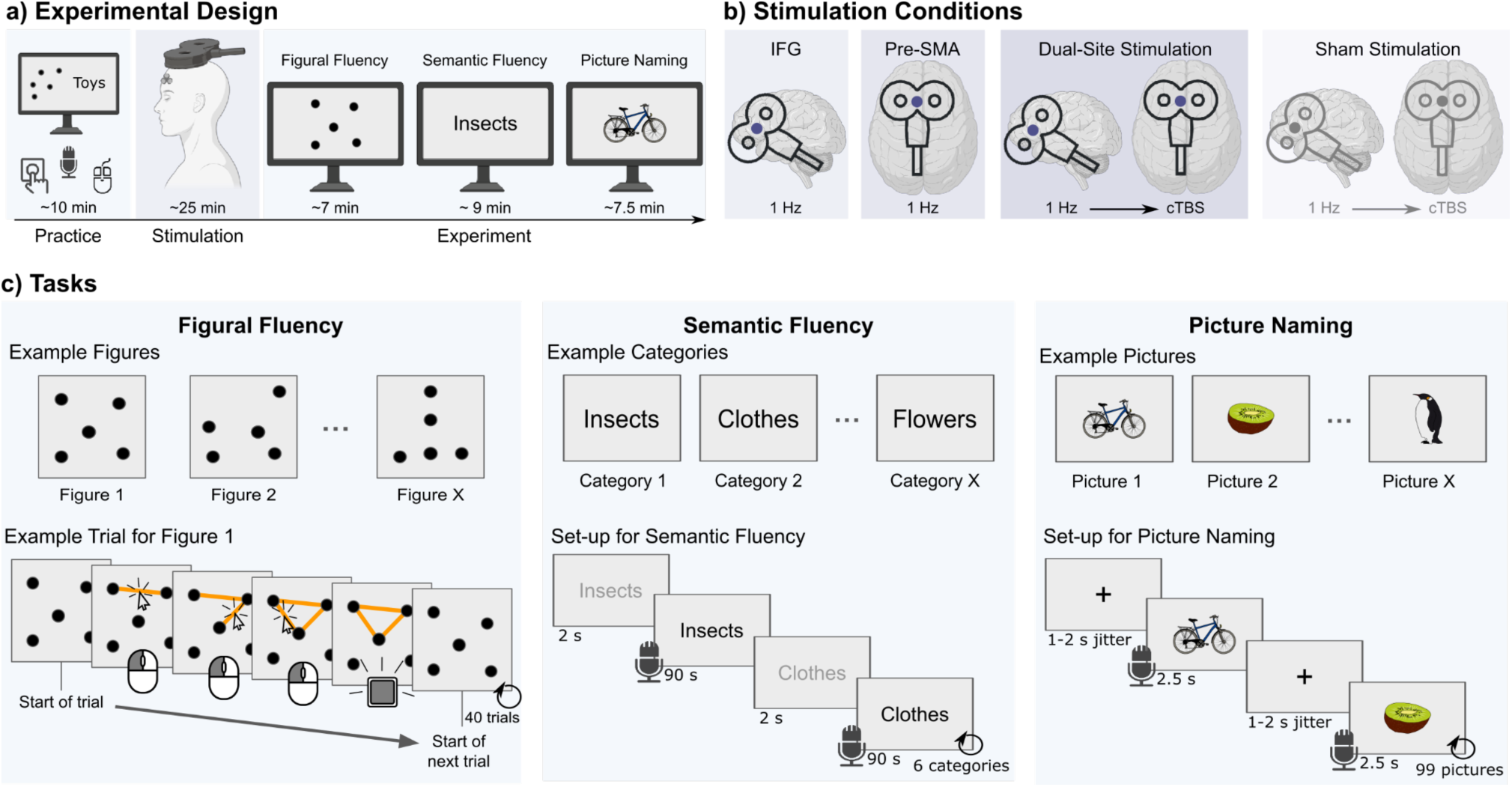
Overview of the study design. a) At the beginning of each session, participants (re-)familiarized themselves with the different tasks. They then received “offline” rTMS and subsequently performed the experiment. The order of figural and semantic fluency in the task battery was counterbalanced across participants. b) All participants received once single-site 1 Hz rTMS to the left IFG and pre-SMA, dual-site rTMS with 1 Hz to the left IFG, succeeded by cTBS to the pre-SMA, and sham rTMS replicating the dual-site condition with a placebo coil. c) During each session, participants completed three blocks of the figural fluency task, six categories of semantic fluency, and one run (99 stimuli) of picture naming.

## Results

We report data from 24 healthy young participants, who each participated in four sessions with rTMS over the left IFG, pre-SMA, dual-site (left IFG succeeded by pre-SMA), and sham stimulation (Fig. 1a and b). After stimulation, participants performed three tasks: verbal semantic and non-verbal figural fluency, and picture naming (Fig. 1c). We were interested in modulatory effects of rTMS on reaction times (RTs) and accuracy in each task.

Following recent methodological considerations for TMS studies in cognition (Ambrosini et al., 2024; Numssen et al., 2023), we included a relatively large sample for a repeated measures design with four sessions and used linear and generalized mixed-effects regression models as well as e-field simulations in our statistical analysis, which allowed us to account for the substantial individual and item-specific variability in response to TMS, while revealing reliable and robust stimulation effects.

### TMS over the left IFG and Pre-SMA both delay Semantic and Figural Fluency

Mixed-effects regression for RTs and accuracy revealed main effects of effective stimulation in all tasks. Specifically, for **semantic fluency**, all stimulation conditions were associated with slower reactions relative to sham stimulation (*β*_IFG_ = 0.09, p = 0.024, *β*_Pre-SMA_ = 0.12, p = 0.007, *β*_Dual site_ = 0.11, p = 0.015, Fig. 2a, Table S1). Stimulation of the pre-SMA induced the relative largest increase in RTs, with an average delay of 189 ms compared to sham. There was no significant difference between the effective stimulation conditions. Results also showed significant main effects for category difficulty with slower reactions for difficult compared to easy categories (*β*_Difficult categories_ = 0.48, p < 0.001, Fig. S2) and slower responses with progressing time (*β*_Trial index_ = 0.002, p = 0.025, Fig. S2). Moreover, there was a significant main effect for executive abilities, where a higher executive score was associated with faster responses (*β*_Executive abilities_ = −0.20, p = 0.028, Fig. S2).

**Figure 2.**
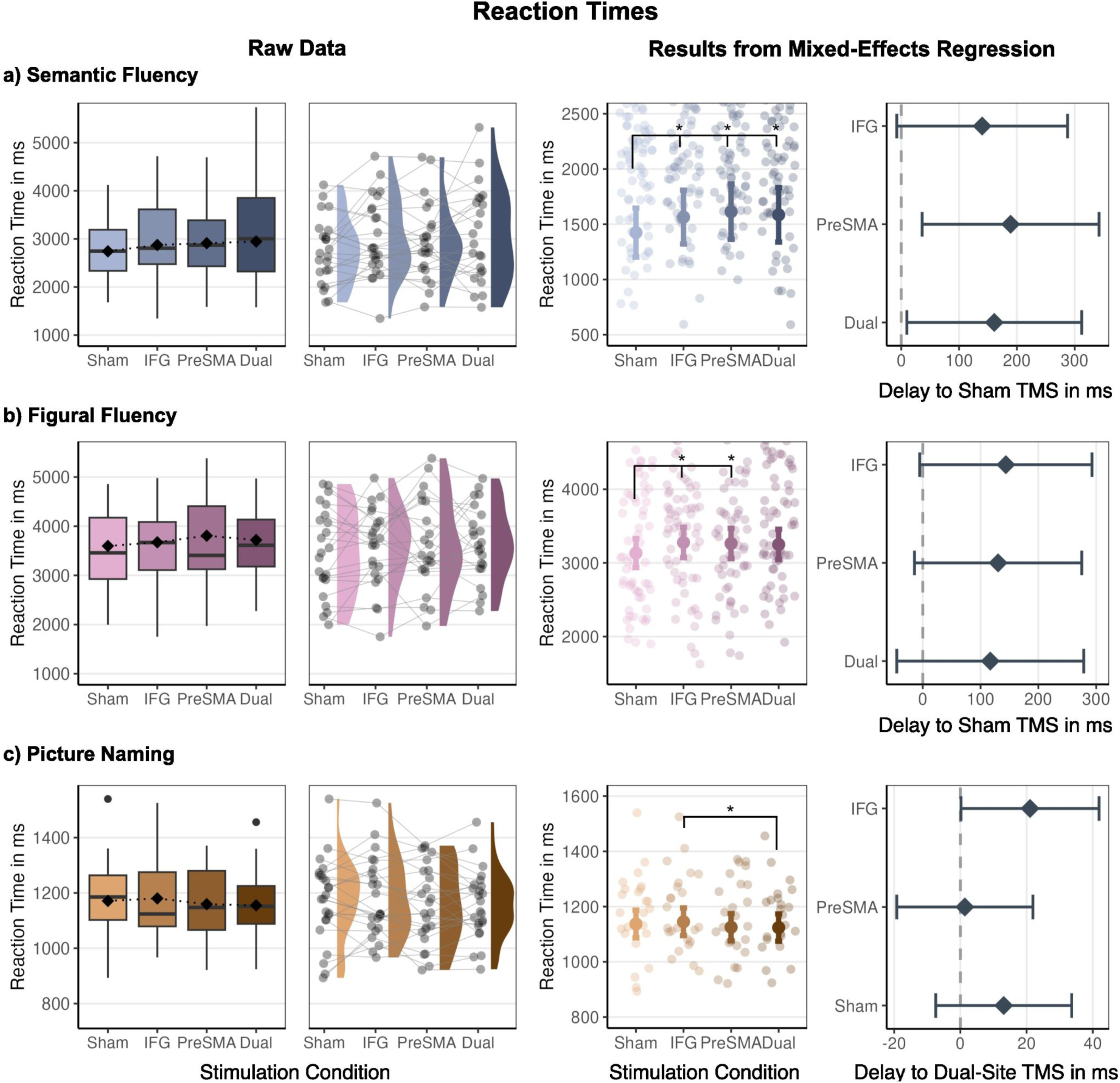
Reaction times: raw data and regression results for each task. Plots on the left side show data aggregated by stimulation condition (boxplots) and distribution of raw data (half-violin plots) with means per participant for each stimulation condition as points. Plots on the right show estimated marginal means from mixed-effects regression models and forest plots displaying the TMS-induced delay in reaction time. Results marked as * p < 0.05 are significant after FDR correction.

Analyzing the number of correct trials per session revealed no significant effect of stimulation for accuracy in semantic fluency (*Incidence rate ratio* (*IRR*)_IFG_ = 0.97, p = 0.506, *IRR*_Pre-SMA_ = 0.96, p = 0.506, *IRR*_Dual site_ = 0.94, p = 0.330, Fig. 3a, Table S2). However, results showed a significant effect of session, such that participants produced more correct items during session 4 compared to session 1 (*IRR*_Session 4_ = 1.10, p = 0.006, Fig. S2). Moreover, there were less correct items for difficult than easy categories (*IRR*_Difficult categories_ = 0.61, p < 0.001, Fig. S2) and higher accuracy with better executive abilities (*IRR*_Executive abilities_ = 1.15, p = 0.003, Fig. S2).

**Figure 3.**
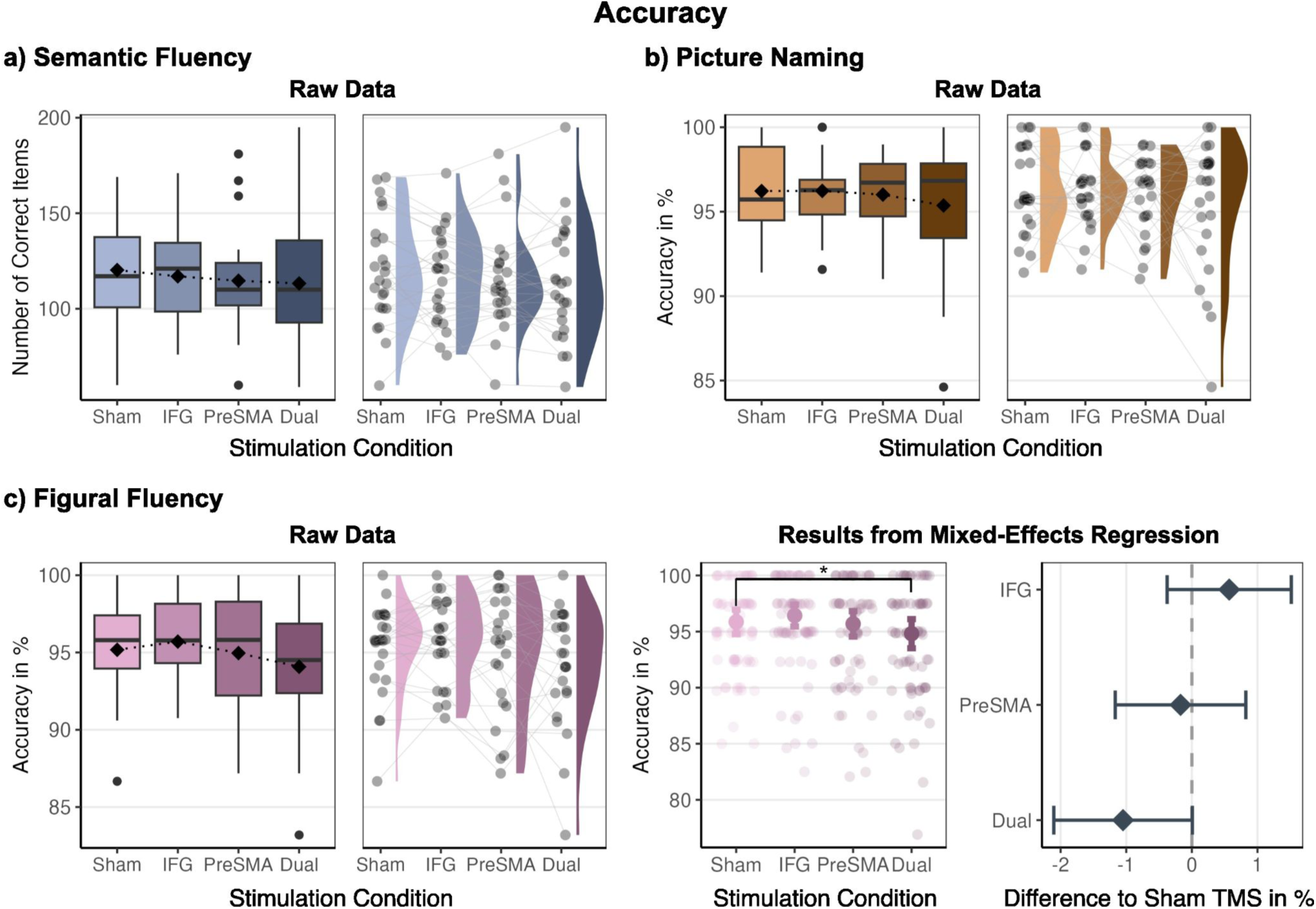
Accuracy: raw data and regression results for each task. **Panels a, b, and left side of c** show data aggregated by stimulation condition (box plots) and distribution of raw data (half-violin plots) with means per participant for each stimulation condition as points. Plots on the **right side of panel c** show estimated marginal means from logistic regression for accuracy in figural fluency and a forest plot displaying the TMS-induced performance decrease. Results marked as * p ≤ 0.05 are significant after FDR correction.

For **figural fluency**, rTMS to both the left IFG and pre-SMA significantly slowed reactions (*β*_IFG_ = 0.04, p = 0.046, *β*_Pre-SMA_ = 0.04, p = 0.046, Fig. 2b, Table S1). The largest effect on RTs in figural fluency was observed for rTMS to the left IFG, with reactions being 144 ms slower compared to sham stimulation. Moreover, results showed a main effect of session, with faster reactions across sessions (*β*_Session 2_ = −0.08, *β*_Session 3_ = −0.14, *β*_Session 4_ = −0.15, all p < 0.001, Fig. S2), indicating a learning effect in this task. Results also revealed significant effects of the number of bars clicked (*β*_N of bars clicked_ = 0.24, p < 0.001, Fig. S2), with slower reactions when participants created more complex designs with more bars activated, and accuracy (*β*_Accuracy_ = −0.32, p = 0.001, Fig. S2), such that higher accuracy was linked to faster reactions. There was also a main effect of executive abilities (*β*_Executive abilities_ = −0.08, p = 0.026, Fig. S2). Similar to semantic fluency, better executive abilities were associated with faster reactions in the figural fluency task.

Analyzing accuracy in the figural fluency task showed worse performance after rTMS to pre-SMA and dual-site compared to sham stimulation (*Odds Ratio* (*OR*)_IFG_ = 1.17, p = 0.239, *OR*_Pre-SMA_ = 0.96, p = 0.736, *OR*_Dual site_ = 0.79, p_uncorr_ = 0.047, p_FDR_ = 0.051, Fig. 3c, Table S2). Moreover, results showed that participants improved their performance in session two compared to session one (*OR*_Session 2_ = 1.45, p = 0.003, Fig. S2) and when they activated a larger number of bars in a figural design (*OR*_N of bars clicked_ = 1.21, p < 0.001, Fig. S2).

Analyzing RTs in **picture naming** showed no significant effect of effective relative to sham rTMS (*β*_IFG_ = 0.01, p = 0.381, *β*_Pre-SMA_ = −0.01, p = 0.203, *β*_Dual site_ = −0.01, p = 0.185, Fig. 2c, Table S1). However, post-hoc tests revealed significantly slower reactions after single-site stimulation to left IFG compared to dual-site stimulation (*β*_IFG - Dual-site rTMS_ = 0.02, p = 0.046), with an average latency of 21 ms. In addition, there was a main effect for trial index with slower reactions with progressing time (*β*_Trial index_ = 0.0003, p = 0.003, Fig. S2) and for executive abilities (*β*_Executive abilities_ = −0.06, p = 0.032, Fig. S2), which reduced reaction times. Similarly, effective rTMS did not impact accuracy in picture naming (*OR*_IFG_ = 1.00, p = 0.989, *OR*_Pre-SMA_ = 0.95, p = 0.905, *β*_Dual site_ = 0.81, p = 0.374, Fig. 3b, Table S2). Results revealed higher accuracy in all sessions relative to session one (*OR*_Session 2_ = 2.09, *OR*_Session 3_ = 1.36, *OR*_Session 4_ = 2.09, all p < 0.05, Fig. S2), as well as with progressing time within a session (*OR*_Trial index_ = 1.01, p = 0.008, Fig. S2), and with higher scores in the STW test (*OR*_STW_ = 1.20, p = 0.035, Fig. S2).

### Task-specific disruption of fluency by stimulation of left IFG and Pre-SMA

To test for a relationship between the induced stimulation strength over the target areas and a change in performance, we correlated individual e-field intensities (Fig. 4a) with mean RTs and accuracy of the respective task and stimulation session. Results showed a significant increase in RTs with stronger e-fields over the left IFG for semantic fluency (r = 0.42, p_FDR_ = 0.039, Fig. 4b). Moreover, linear regression combining both e-fields for sessions with dual-site stimulation showed poorer performance in figural fluency with increasing intensity of the e-field in pre-SMA (r = −0.55, p = 0.043, Fig. 4b). Fig. S3 shows results for all correlations.

**Figure 4.**
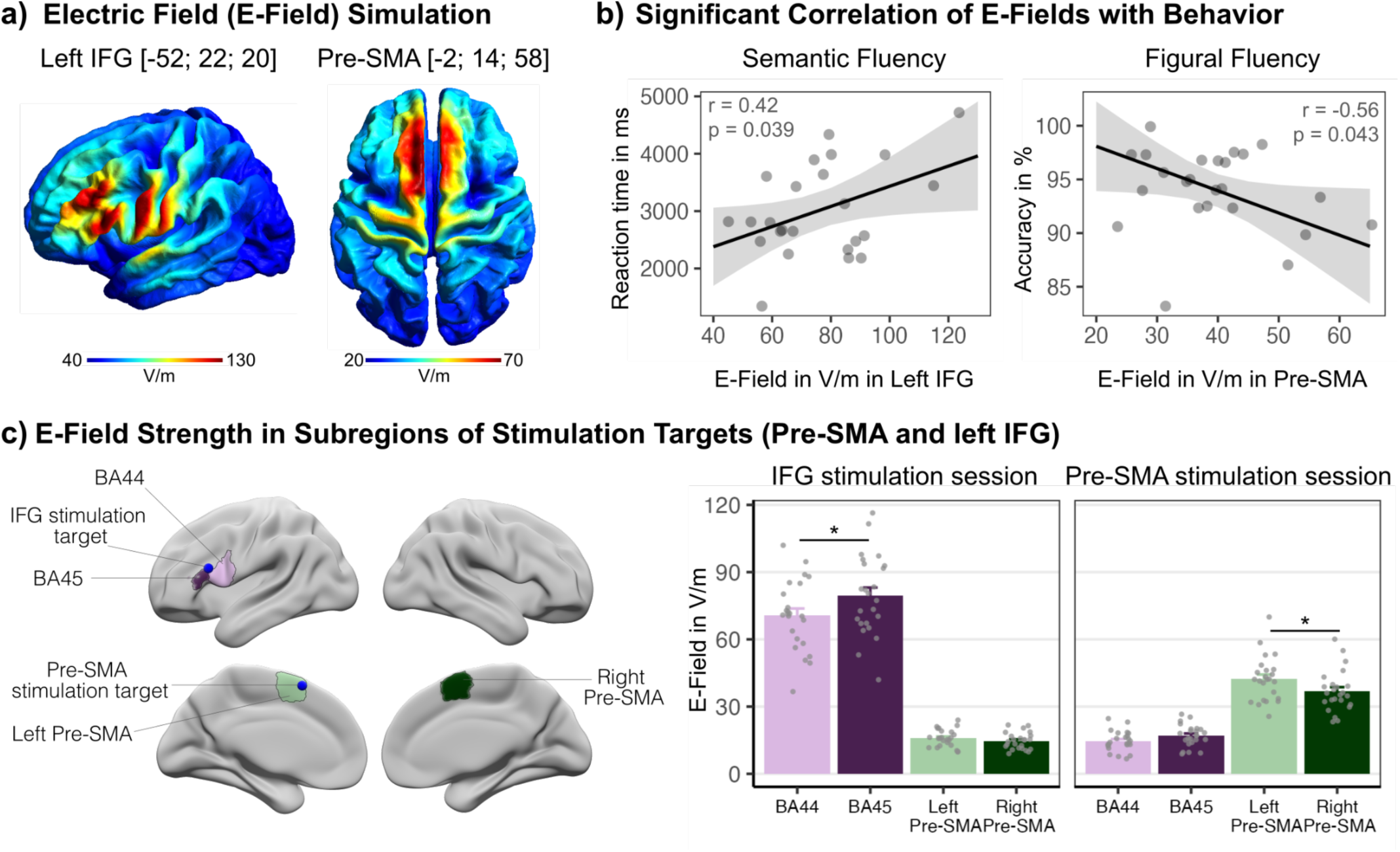
Task-specific performance disruptions induced by the electric field (e-field) strength. **Panel a** shows the induced e-fields over our stimulation targets in the left posterior IFG and pre-SMA. **Panel b** displays significant correlations of e-fields with behavior after FDR correction. Stronger e-fields in the IFG were associated with slower reactions during semantic fluency, while stronger e-fields in the pre-SMA were linked to poorer performance in figural fluency. **Panel c** shows the e-field strength in subregions of our stimulation targets, anterior (BA45) and posterior (BA44) IFG, and left and right pre-SMA. Anatomical parcels are based on cytoarchitectonic probabilistic maps (Julich-Brain Atlas).

We were interested in the specific strength of the TMS effect in subregions of each stimulated cortical area and extracted e-field values in cytoarchitectonically defined ROIs. Comparing e-field strengths between ROIs and stimulation sessions revealed a significant interaction between both predictors (*X* = 1648.44, p < 0.001), with higher e-field values in IFG and pre-SMA ROIs in the respective stimulation session (Fig. 4c, Table S3). Furthermore, in the IFG session, the anterior IFG (BA45) received significantly more stimulation than the posterior IFG (BA44; *β*_BA45 - BA44_ = 8.87, p < 0.001) and during rTMS over pre-SMA, the left pre-SMA showed stronger stimulation strength than the right pre-SMA (*β*_left pre-SMA - right pre-SMA_ = 5.47, p = 0.011).

### TMS significantly impairs strategic processes in Semantic Fluency

TMS may induce behavioral changes extending beyond accuracy and reaction times, potentially altering task strategies. We investigated whether rTMS influenced clustering and switching, two core processes for cognitive flexibility in semantic fluency, by comparing the number of switches between stimulation conditions. Results showed a significant effect of pre-SMA and dual-site rTMS but not IFG rTMS (*IRR*_Pre-SMA_ = 0.90, p = 0.039, *IRR*_Dual site_ = 0.87, p = 0.007, *IRR*_IFG_ = 0.93, p = 0.143, Fig. 5, Table S4). Both pre-SMA and dual-site rTMS reduced the number of switches compared to sham stimulation. The most pronounced effect was observed with dual-site rTMS, which resulted in an average reduction of 1.25 switches compared to sham stimulation. Moreover, there was an effect of category difficulty with less switches for difficult than easy categories (*IRR*_Difficult categories_ = 0.60, p < 0.001), but no effect of session (all p > 0.05, Table S4). Figure S4 shows the average number of switches per semantic category.

**Figure 5.**
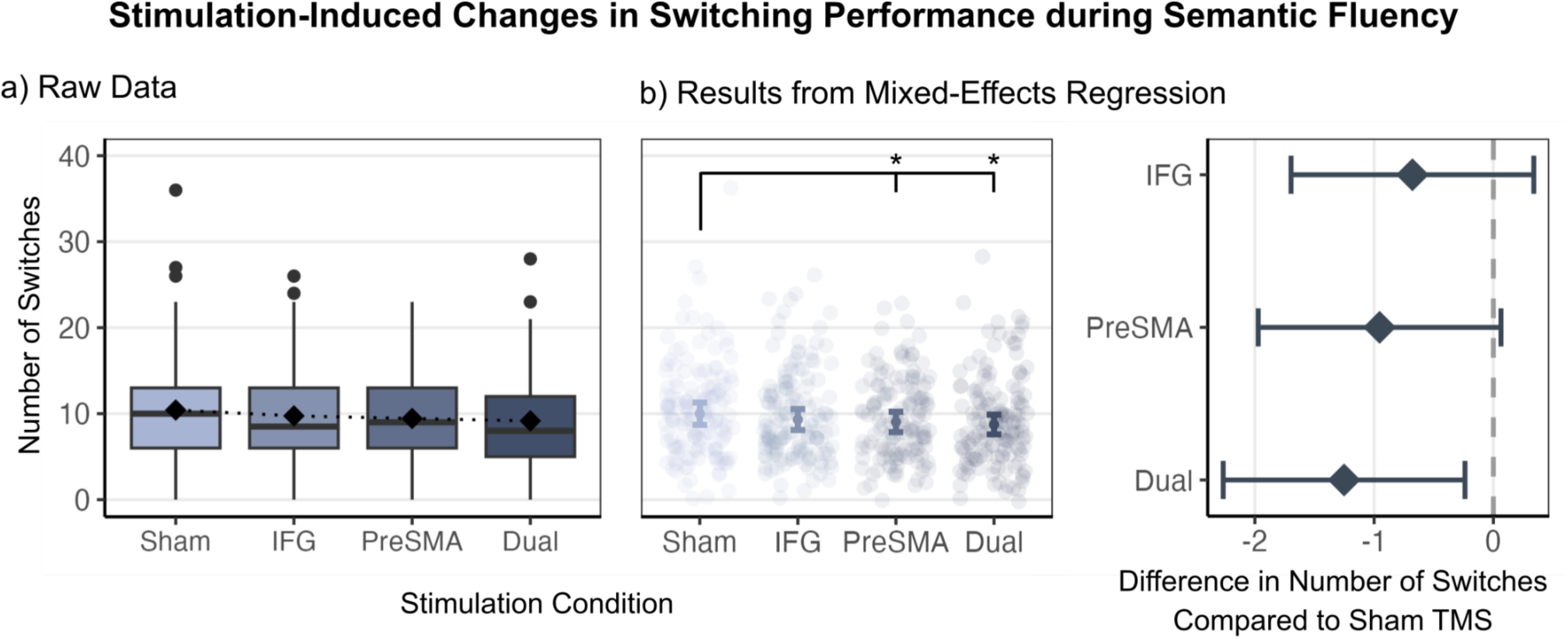
Stimulation-induced changes in the number of switches during semantic fluency. Both pre-SMA and dual-site rTMS disrupt semantic fluency through fewer switches within semantic categories. Results marked as * p < 0.05 are significant after FDR correction.

## Discussion

This study provides new insight into the functional relevance of the left IFG and pre-SMA in semantic-specific and domain-general executive control. While stimulation of either region broadly disrupted both semantic and figural fluency, suggesting some shared functionality, we identified distinct specializations through the analysis of stimulation strength and behavioral outcomes. The left IFG was primarily linked to semantic control, as evidenced by its association with verbal fluency deficits, whereas the pre-SMA was more involved in domain-general executive control, impacting non-verbal fluency and cognitive flexibility, such as clustering and switching during semantic fluency. Additionally, dual-site TMS targeting both the IFG and pre-SMA, delayed semantic fluency and impaired accuracy in figural fluency. Notably, the impaired accuracy in figural fluency was only observed after dual-site stimulation, providing first evidence for successful compensation of executive functions through either the left IFG or pre-SMA following perturbation of the other area. Overall, these findings stress the multidimensionality of cognitive control. They support a hierarchical model where semantic tasks recruit both specialized (IFG-mediated) and general executive (pre-SMA-mediated) resources, and demonstrate that executive control is based on a closely interacting network subserved by the left IFG and pre-SMA.

Contrary to our hypothesis, perturbation of the IFG did not only impair semantic but also figural fluency, indicating a functional role of this area in a non-verbal, strongly executive task. This finding aligns with a growing body of research suggesting the involvement of the IFG in multiple cognitive domains beyond language and semantics (Friederici, 2011; Klaus & Hartwigsen, 2019; Lambon Ralph et al., 2017), including social cognition (Diveica et al., 2021) and executive control (Fedorenko et al., 2012; Fedorenko & Blank, 2020). However, what remains unclear is whether these different functions are reflected by distinct subregions within this cortical area or whether they show at least some functional overlap. Our target coordinate in the IFG was located in the dorsal anterior IFG (corresponding to Brodmann area 45), which has been repeatedly associated with semantic processing based on functional activity and connectivity (Klaus & Hartwigsen, 2019; Nieberlein et al., 2024; van der Burght et al., 2023; Wagner et al., 2014). The observed effect on figural fluency might be due to several reasons. First, recent investigations revealed a functional organization of the IFG along gradient axes where the dorsal IFG has been attributed a domain-general function of executive control (Chiou et al., 2023; Diveica et al., 2023), aligning with previous observations that the more language-specific anterior IFG is neighbored by a domain-general posterior region (Fedorenko et al., 2012).

Second, our target coordinate was proximal to the posterior IFG (corresponding to Brodmann area 44). Although functional divisions do not strictly follow cytoarchitectonic borders, the stimulation likely affected this more domain-general region, influencing both fluency tasks. This aligns with van der Burght et al. (2023), who demonstrated the challenge of dissociating TMS effects between anterior and posterior regions in the left IFG. Our e-field analysis of subregions revealed that while stimulation was statistically stronger in the anterior IFG, the posterior region still received substantial stimulation (71 V/m on average). To address this lack of focality, we calculated the effective induced stimulation strength, which is shaped by individual anatomical characteristics, at our target sites (Numssen et al., 2023). Crucially, this analysis unveiled a domain specificity for each region: only TMS over the IFG significantly impacted verbal fluency. This finding suggests that despite potential stimulation spread, the anterior IFG maintains a primary role in semantic control.

The effect of pre-SMA stimulation on both fluency tasks aligned with our expectations and confirms its domain-generality. The pre-SMA is considered a hub of the MDN (Camilleri et al., 2018; Duncan, 2010), critical for response inhibition, cognitive flexibility, and conflict monitoring (Iannaccone et al., 2015; Nachev et al., 2008; Obeso et al., 2013). More recently, it has also been linked to semantic control processes in neuroimaging (Chiou et al., 2023; Jackson, 2021) and neurostimulation (Martin, Frieling, et al., 2023) research, as well as to compensatory roles in language processing in the aging brain (Martin et al., 2022; Martin, Williams, et al., 2023) and post-stroke aphasia (Geranmayeh et al., 2017; Ren et al., 2023). Importantly, our findings support an overall executive aspect of this region, which exhibits broad domain generality and involvement in executive control across domains. This conclusion is strengthened by the impaired accuracy in figural fluency with increasing pre-SMA stimulation and the reduced number of switches in semantic fluency following TMS to the pre-SMA. Switching between subcategories in semantic fluency requires cognitive flexibility and has been identified as an indicator of the successful search process during semantic fluency (Lundin et al., 2023; Troyer et al., 1997) and a detector of cognitive impairments (Nikolai et al., 2018; Troyer et al., 1998). Our results confirm the strongly executive nature of fluency tasks (Aita et al., 2019; Ghanavati et al., 2019; Shao, 2014). Moreover, they reconcile the role of the pre-SMA in semantic control with its overarching function in coordinating distributed executive resources during complex tasks.

We applied dual-site TMS, targeting the IFG followed by the pre-SMA, to test whether disrupting both areas would result in greater fluency deficits compared to single-site stimulation (Bergmann & Hartwigsen, 2021). While no additional effect of dual-site TMS was observed for semantic fluency, figural fluency accuracy was impaired exclusively after dual-site stimulation. E-field analysis revealed that this effect was linked to the stimulation strength in the pre-SMA. This finding further underscores the key role of the IFG in domain-general executive control (Dick et al., 2019) and aligns with the delayed reactions in figural fluency after single-site TMS to both the IFG and pre-SMA. It is particularly noteworthy that we only observed this effect after dual-site TMS, suggesting that when either site is perturbed individually, successful compensatory mechanisms may mitigate the impact of TMS, at least for response accuracy. The fact that response efficiency was already affected after uni-site stimulation over either area converges with previous reports that response speed may be generally more sensitive to stimulation-induced perturbations than accuracy (Bergmann & Hartwigsen, 2021; Bonaiuto et al., 2016). The significant impact of dual-site TMS on figural fluency accuracy highlights the importance of both regions in executive control, while the absence of this effect in semantic fluency further strengthens the unique capacity of the IFG in both domain-specific semantic and domain-general executive control.

Finally, we used picture naming as a low-level task which requires little cognitive control in healthy young adults but, like semantic fluency, engages targeted lexicosemantic retrieval. Consistent with this expectation, we observed delayed reactions in picture naming after TMS to the IFG, supporting its established role in naming processes (Ferré et al., 2020; Jarret et al., 2022; Xing et al., 2018). No other stimulation-induced effects were observed for this task, likely due to participants’ fast and highly accurate performance and the experimental design, where picture naming was always presented last, diminishing the temporary TMS effect.

In summary, this study probed the functional relevance of the left IFG and pre-SMA in semantic and executive control by inhibiting either site alone or together using rTMS. Examining the impact of these disruptions on verbal and non-verbal fluency tasks revealed significant contributions of both regions to domain-specific and domain-general control processes. The left IFG was found to be crucial for semantic control, particularly in verbal fluency tasks, while the pre-SMA played a more domain-general role in executive functions, affecting both semantic and non-semantic tasks. Notably, only dual-site stimulation impaired accuracy in figural fluency, suggesting compensatory mechanisms between these regions in executive control that help to maintain accurate responses even if response efficiency is reduced as evidenced by delayed response speed after uni-site stimulation. Our findings support a hierarchical model of cognitive control where semantic tasks engage both specialized and general executive resources. Furthermore, they provide evidence for a network of executive control subserved by the left IFG and pre-SMA, contributing to a deeper understanding of the neural basis of semantic and executive functions.

## Materials and Methods

### Participants

We tested 24 healthy adults (M = 30.00, SD = 5.32, range: 20–40 years, 13 female), who were right-handed, native German speakers, and had no contraindication to MRI and TMS. Vocabulary and verbal intelligence were assessed with the German version of the Spot-the-Word test (STW; Schmidt & Metzler, 1992) and general executive functions with the Trail Making Test (TMT; Reitan, 1958) and the Digit Symbol Substitution Test (DSST; Wechsler, 1944). Participants were informed about experimental procedures but blinded to the different TMS conditions and their order. The study was approved by the local ethics committee of the Medical Faculty at Leipzig University and conducted in accordance with the Declaration of Helsinki. Written informed consent was obtained from each participant prior to the experiment.

### Design

We employed a repeated-measures within-subjects design with four sessions per participant, which were separated by at least one week. At the beginning of each session, participants (re-)familiarized themselves with the task. They then received offline rTMS stimulation and subsequently performed the experiment (Fig. 1A). The study comprised four stimulation conditions: IFG, pre-SMA, dual TMS (IFG first, followed by pre-SMA stimulation), and sham stimulation (Fig. 1B). The order of these conditions was counterbalanced across subjects.

### Tasks and Stimuli

Participants performed three tasks after offline rTMS: semantic fluency, figural fluency, and picture naming. The experiment always began with both fluency tasks to ensure maximal post-stimulation effects. Half of the participants started with semantic fluency, followed by figural fluency, and vice-versa for the others.

The semantic fluency task consisted of 24 semantic categories, containing easy (e.g., clothes) and difficult (e.g., insects) items (see Fig. S4 for all categories). Difficulty ratings were taken from a previous study, for which we piloted difficulty levels of semantic categories (Martin et al., 2022). Participants completed six categories during one session, consisting of three easy and three difficult categories, pseudorandomized in their order. For each category, participants were instructed to name as many unique items as possible during 90 s, avoiding repetitions and proper names (Fig. 1C).

We employed the Five-Points Test (5PT; Regard et al., 1982), a standardized assessment of figural fluency (Tucha et al., 2012), as a non-verbal measure of fluid intelligence. This test requires participants to generate abstract geometrical figures within a time limit, imposing executive demands similar to the semantic fluency task. We adopted a computer-based version of the 5PT, which we piloted online in a separate sample of 30 young adults (M = 23.5, SD = 23.7, range: 18–30 years). To this end, we developed eleven additional 5-dot arrangements as stimuli (Fig. S1). Although the dot arrangements differed in geometrical regularity and internal symmetry to the original design of the 5PT, we controlled for dimensions on screen and ease of clickability by matching the cumulative bar length of each figure. Participants completed 3 different dot arrangements per session, each lasting a maximum of 3.5 min or until 40 designs were completed, and were instructed to produce as many unique designs as possible by connecting two or more dots with straight lines (Fig. 1C).

The picture-naming task was implemented as a low-level control task, accounting for bottom-up access processes implicated in word retrieval. We compiled four stimulus lists with 99 pictures each. Pictures were taken from the Multipic database (Duñabeitia et al., 2018) and balanced for naming agreement, visual complexity, word frequency, and word length in syllables. Each picture was presented for 2.5 s, with a total duration of 7.5 min for picture naming per session (Fig. 1C). All tasks were programmed and presented with Psychopy version 2021.2.3.

### Transcranial Magnetic Stimulation

Before the experimental tasks, we applied 1 Hz offline rTMS at 100% resting motor threshold (rMT) for 25 min over either the left IFG or pre-SMA. During the session with dual-site stimulation, 1 Hz rTMS to the left IFG was immediately succeeded by continuous theta burst stimulation (cTBS) to pre-SMA. cTBS was delivered at 90% rMT (cf. Williams et al., 2024) and consisted of 600 pulses delivered in 50 Hz triplets every 200 ms for a total of 40 s. We chose to employ cTBS as second stimulation protocol during dual-site TMS to avoid extended stimulation time during this session. To maximize effective blinding, the sham session mimicked the dual-site stimulation using a placebo coil.

The rMT was assessed at the beginning of the first session and defined as the lowest stimulation intensity inducing motor evoked potentials of ≥ 50 µV at least 5/10 times in the relaxed first dorsal interosseous muscle when single-pulse TMS was applied to the left motor cortex. TMS was delivered with a figure-of-eight coil (MagVenture MCF-B65 for effective stimulation and MCF-P-B65 placebo coil for sham stimulation) connected to a MagPro X100 stimulator (MagVenture) and guided by stereotactic neuronavigation (TMS Navigator, Localite). To this end, the participant’s head was co-registered onto an individual structural MR scan taken from the in-house database or newly acquired at a 3T Siemens MR scanner. The coil was oriented at 0° for pre-SMA (Casula et al., 2022) and at 45° for IFG stimulation (Fig. 1B). Target locations for the left IFG (x: −52, y: 22, z: 20) and pre-SMA (x: −2, y: 14, z: 58) were derived from a meta-analysis on activation peaks in semantic fluency tasks (Wagner et al., 2014). The MNI coordinates were converted into participants’ native space and used for neuronavigation.

We performed post-hoc e-field simulations using SimNIBS v.4.0.0 (Thielscher et al., 2015) to characterize location, extent, and strength of the e-field induced by rTMS over the pre-SMA and IFG in each participant. E-field values were extracted by defining a region of interest with a 5 mm radius around the stimulation coordinates. From this region, values at the 95% percentile were extracted from the individually calculated e-fields, focusing exclusively on gray matter. To assess the relationship between the induced stimulation strength over the target areas and a change in performance, we correlated the individual by-target-site induced e-field with the individual mean reaction time (RT) and accuracy per task and session. For the session with dual-site stimulation, we calculated linear regression models using individual e-fields from both targets as predictors and behavioral performance as outcome measure.

To further understand how the stimulation affected subregions in IFG and pre-SMA, we additionally extracted e-field values within cytoarchitectonic probabilistic regions of interest (ROI) in both cortical areas taken from the Julich-Brain Atlas (K. Amunts et al., 2021). Probabilistic maps of anterior (BA45) and posterior IFG (BA44), and left and right pre-SMA were thresholded at 35%, binarized, and resampled to subject space, before extracting the 95% quantile in each region. A linear mixed-effects model with ROI and session (stimulation target IFG or pre-SMA) as fixed effects and by-participant random intercepts was used to compare e-field values in the ROIs.

### Data Analysis

#### Preprocessing

Recordings from the semantic fluency and the picture naming tasks were transcribed by three native German raters who annotated reaction times with Praat (Boersma & van Heuven, 2001) and attributed accuracy scores to each response. For semantic fluency, repetitions and wrong answers were marked as incorrect. For picture naming, answers that did not match with the names prescribed by Multipic’s norms were evaluated by the raters. For figural fluency, repeated designs were marked as incorrect.

We obtained RTs for each task according to the following criteria: For semantic fluency, we extracted the interval between the offset of the previous item and the onset of the following item. For figural fluency, RTs were calculated as the interval between the presentation of a blank set of dots and the subject’s button press to record their design, resulting in a maximum of 40 trials per figure. For picture naming, we considered the interval between picture presentation and naming onset. To exclude excessive outlier values from RTs, we employed a lenient approach where for each participant, session, task, and stimulus item (category, figure), RTs at least three SDs above the mean were excluded. This procedure removed 3.6% of all trials.

#### Statistical Analysis

For each task, accuracy and RTs of each session were analyzed using mixed-effects models. Linear regression models were applied to log-transformed RTs, considering only correct trials. Accuracy data were analyzed using logistic regression for the binomially distributed data of figural fluency and picture naming, and negative binomial regression for the count data of semantic fluency.

Table 1 presents the task-specific models for reaction time and accuracy. Based on our research question, stimulation condition (i.e., sham, pre-SMA, IFG, or dual-site rTMS) was always included as a fixed effect. Additionally, all models incorporated fixed effects for session to account for potential learning effects and trial index to account for the linearly decreasing TMS effect during a session. Moreover, we included scores from neuropsychological tests as measures of verbal intelligence and executive abilities. Both executive tests (TMT, DSST) were combined into a single score. This was done by first inverting the reaction times of the TMT, then z-standardizing both the TMT and DSST, and finally calculating the individual mean of both tests. Scores from the Spot-the-Word test were z-standardized and included as a separate regressor, as it assesses verbal knowledge without time restrictions and thus represents a different cognitive concept. Finally, each model included task-specific predictors, such as category difficulty for semantic fluency and number of bars clicked for figural fluency.

**Table 1.**
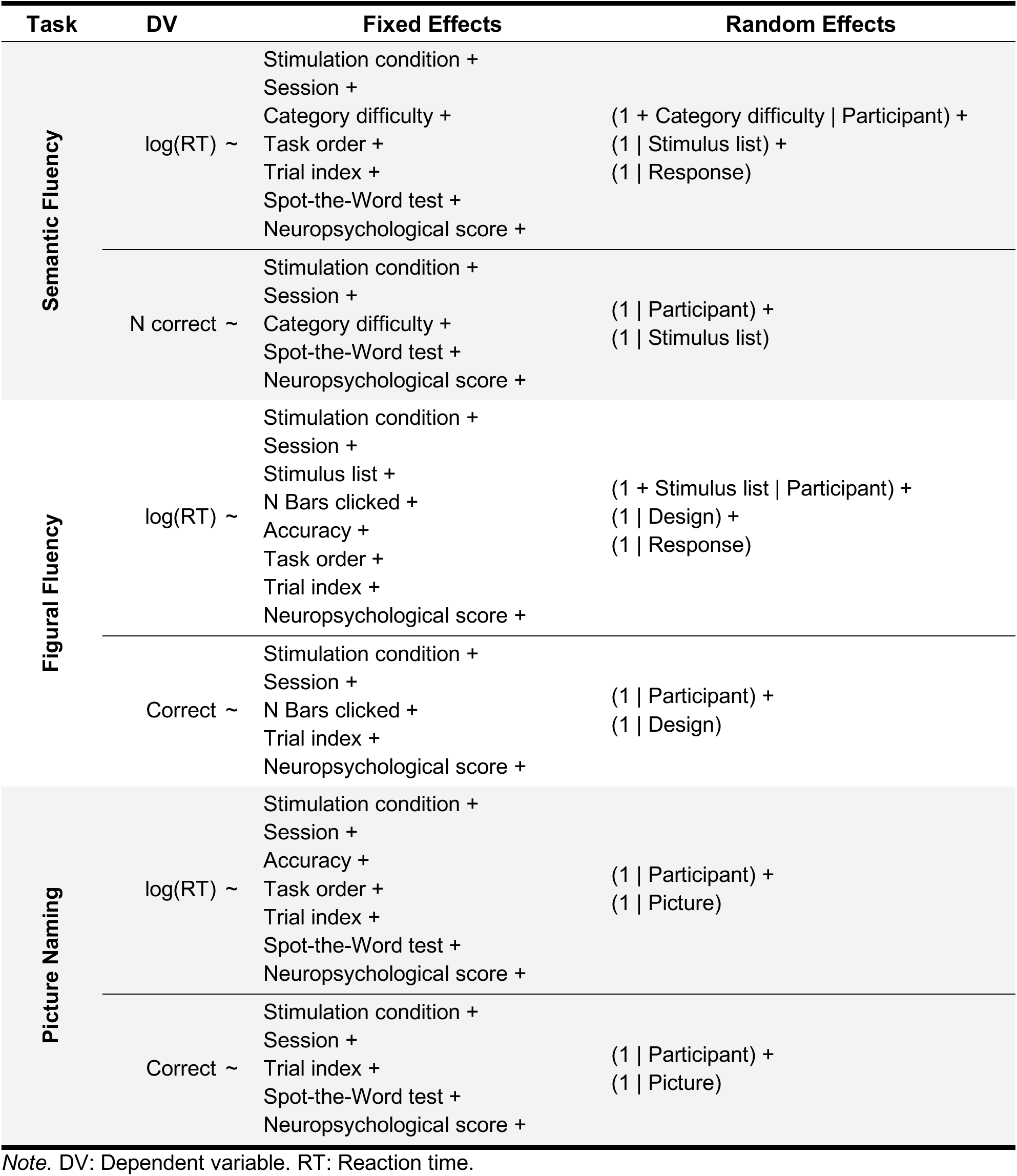
Mixed-effects models for reaction times and accuracy.

Stepwise model selection was employed to determine the best-fitting model based on the Akaike Information Criterion (AIC) and model convergence, with a change of at least two points in AIC considered meaningful (Burnham & Anderson, 2004). All models included at least two random intercepts to account for variance introduced by participants and stimuli. Where possible, RT models also included task-specific by-participant random slopes. All factors were contrast coded using the simple coding scheme, with *sham stimulation* and *session 1* as reference levels.

Statistical models were conducted using R v.4.4.1 (R Core Team, 2024) with the lme4 package (Bates et al., 2015) for mixed-effects models and the performance package (Lüdecke et al., 2021) for model comparisons. Post-hoc multiple comparisons were performed using the emmeans package (Lenth, 2020) to determine effects between individual factor levels other than the reference levels, with FDR correction applied. Plots were generated using ggplot2 (Wickham, 2016) and ggeffects (Lüdecke, 2018), and model output via sjPlot (Lüdecke, 2021). All reported p-values for stimulation condition are FDR-corrected to account for comparisons of each condition to sham in the output of the mixed-effects models.

#### Exploratory quantitative analysis for semantic fluency

To further investigate the effects of TMS on the executive components of verbal fluency, we conducted an exploratory quantitative analysis on clustering and switching strategies within the verbal fluency task. To evaluate participants’ lexical retrieval, we employed FastText (Bojanowski et al., 2017) for fully automated switch detection. This method estimates the number of switches among clusters by calculating the cosine similarity between the vector of any uttered word and the subsequent word vector. If the cosine similarity falls below a predefined switch threshold, a new cluster is identified. We defined category-specific thresholds as the median similarity across all word pairs within a given semantic category as suggested by Alacam et al. (2022). We hypothesized that effective stimulation would impact lexico-semantic retrieval as well as clustering and switching processes, two core strategies underlying fluency performance (Troyer et al., 1997), thus resulting in fewer switches. To test this hypothesis, we ran a negative binomial mixed-effects regression with the number of switches per participant and category as dependent variable and stimulation condition, session, and category difficulty as fixed effects. Moreover, we included random intercepts for participants and categories.

## Data Availability

Data and analysis scripts are available in our OSF repository https://osf.io/q5kam/.

## Acknowledgments

We would like to thank Julia Siodmiak for assistance with data acquisition. Moreover, we thank all student assistants and interns involved in transcriptions of verbal fluency and picture naming data. We thank Ole Numssen for advice with the calculations of the electric fields.

## Author contributions

SM: Conceptualization, data curation, formal analysis, investigation, methodology, project administration, visualisation, writing: original draft, review & editing

MF: Conceptualization, data curation, formal analysis, investigation, methodology, project administration

AB: Formal analysis, methodology, writing: review & editing

GH: Conceptualization, funding acquisition, supervision, methodology, writing: review & editing

## Supplementary Materials

**Figure S1.**
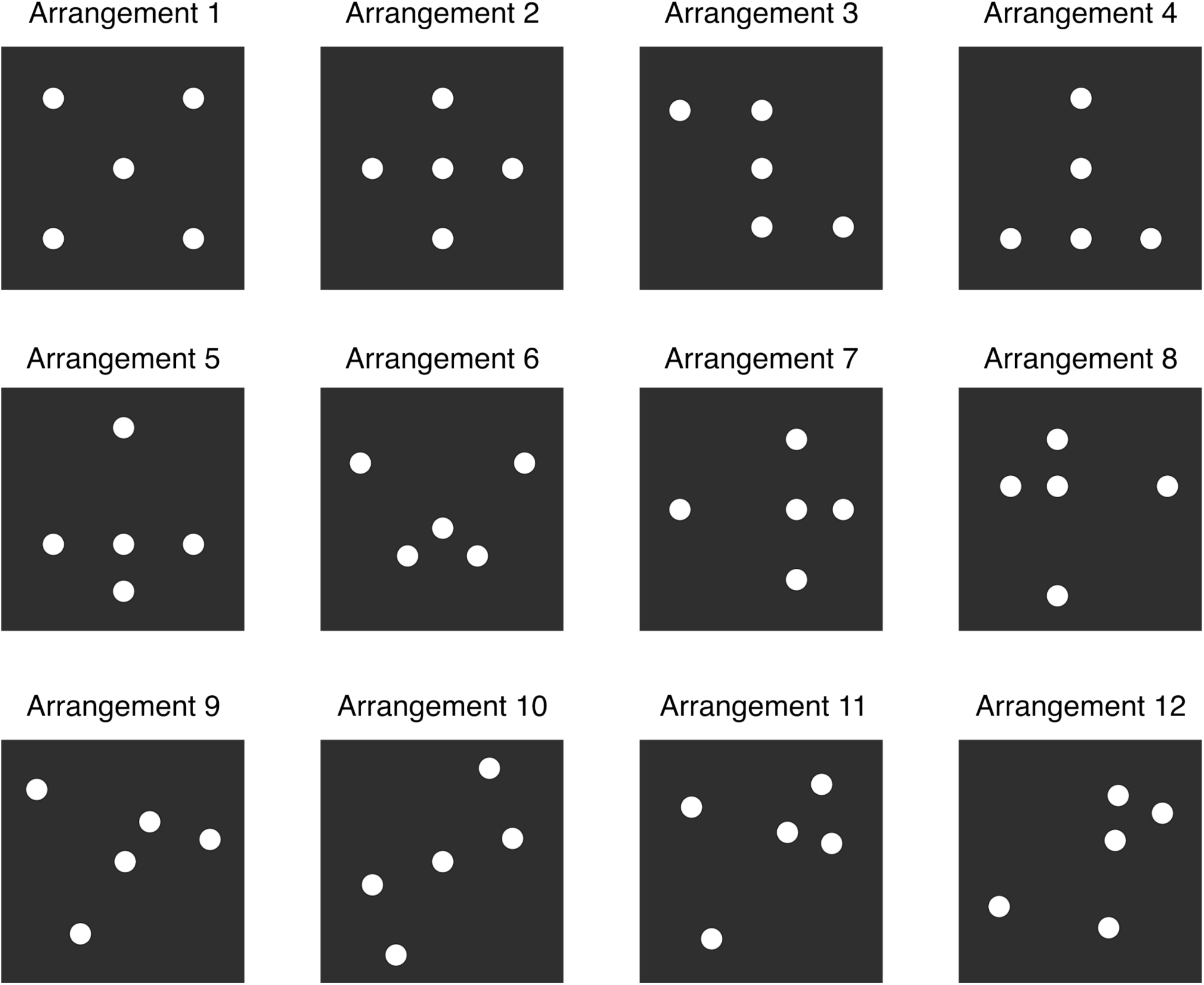
Dot configurations for five-point task. We developed a computerized version of the five-point task and designed 11 additional dot arrangements so that participants had to complete three unique dot arrangements per session.

**Figure S2.**
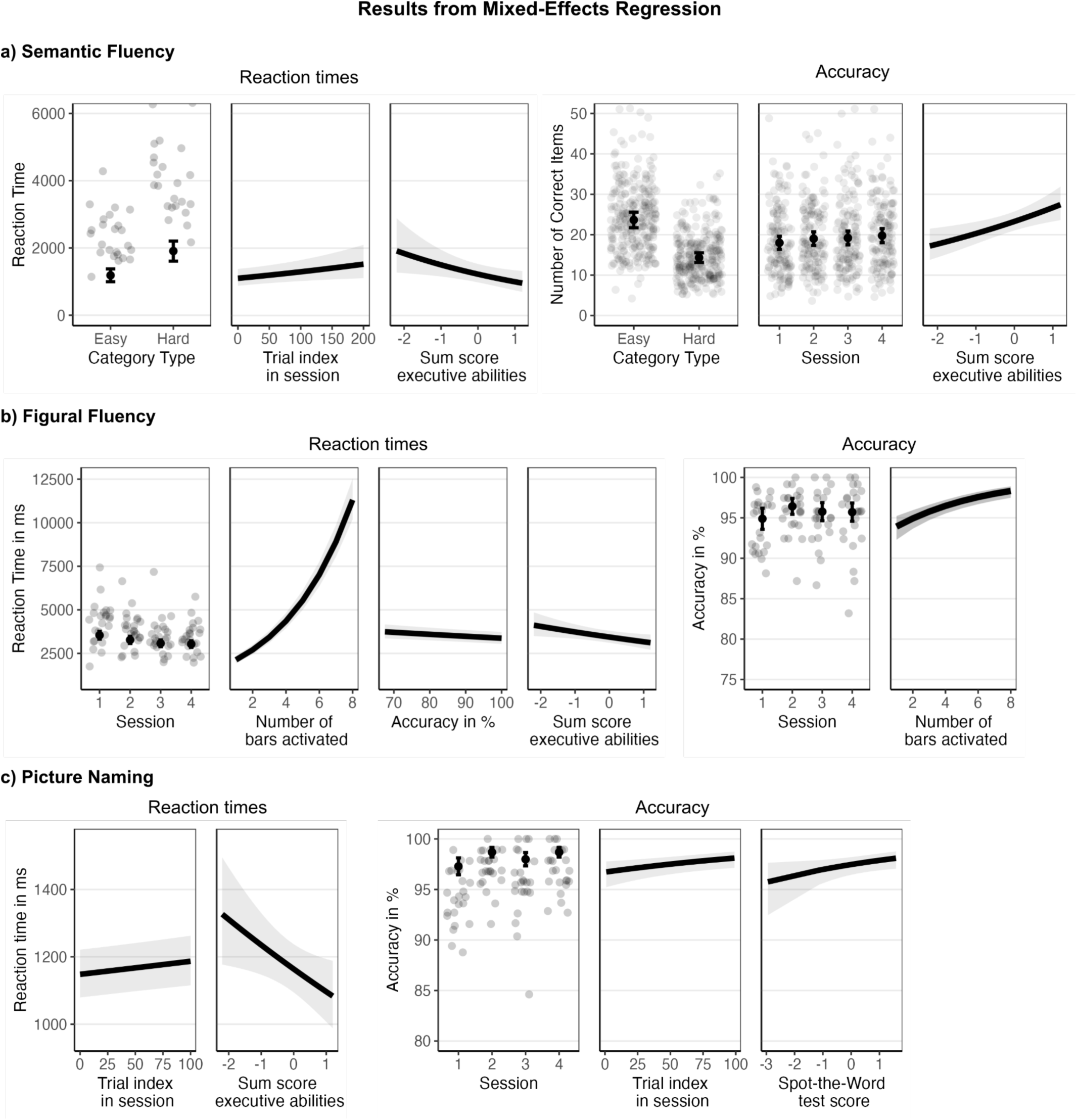
Results from mixed-effects regression. Panels show significant main effects for other predictors than stimulation condition for reaction times and accuracy in each task.

**Table S1.**
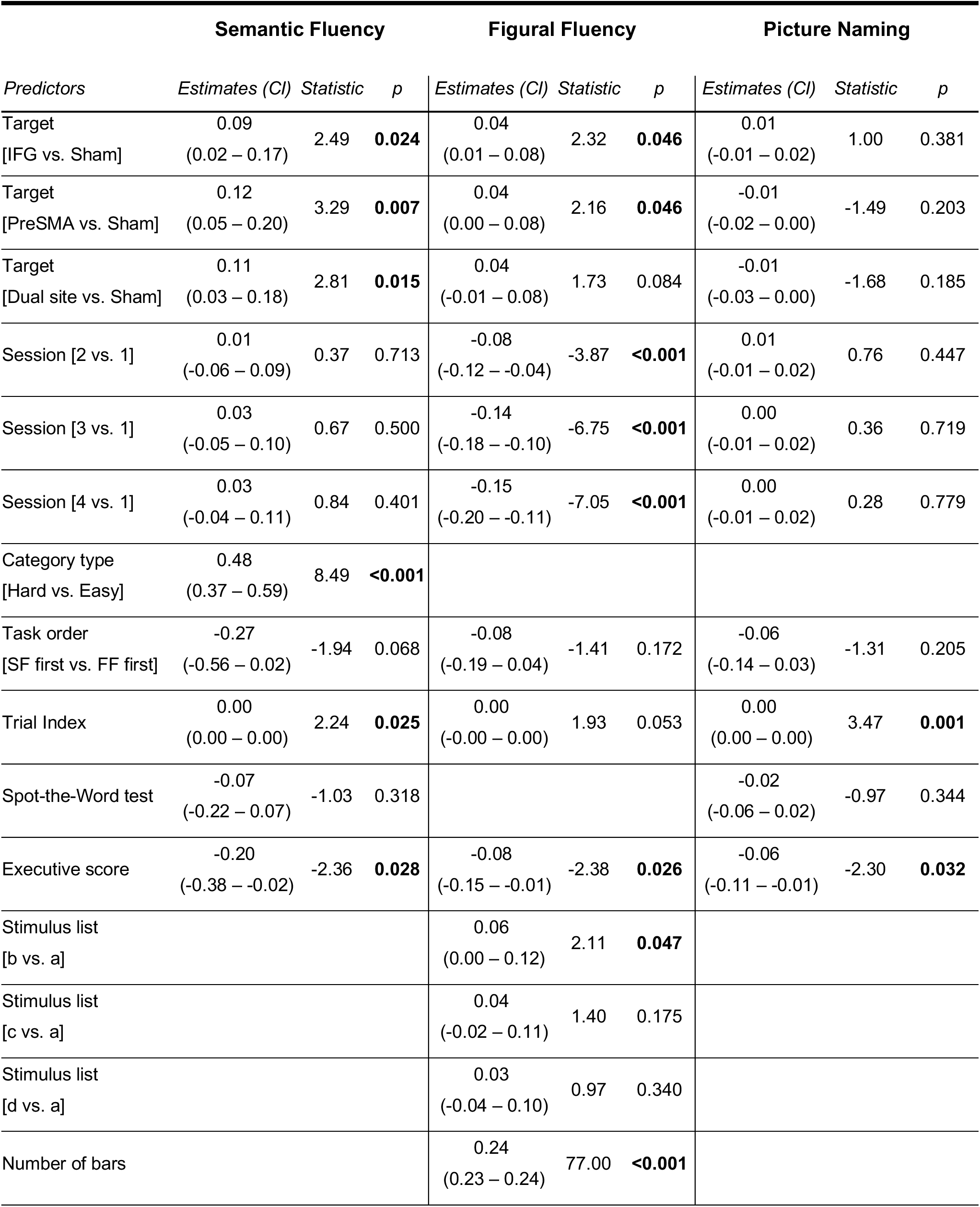
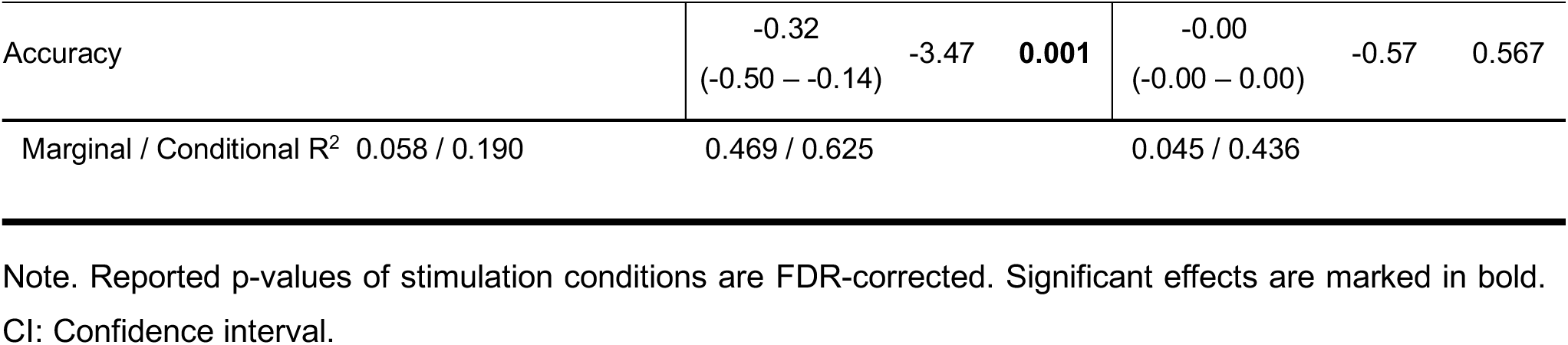
Reaction times: Results from mixed-effects regression for each task.

**Table S2.**
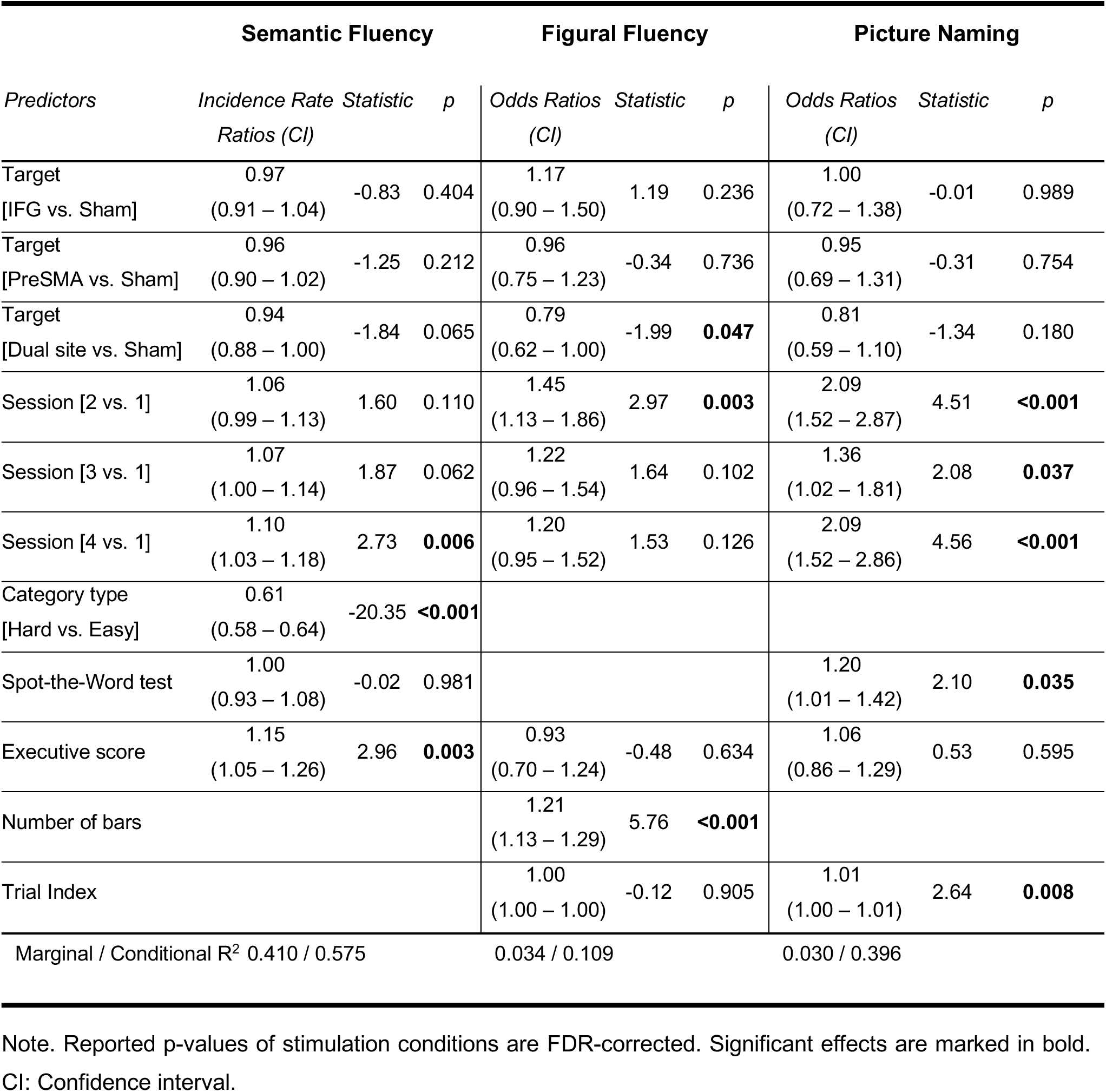
Accuracy: Results from mixed-effects regression for each task.

**Figure S3.**
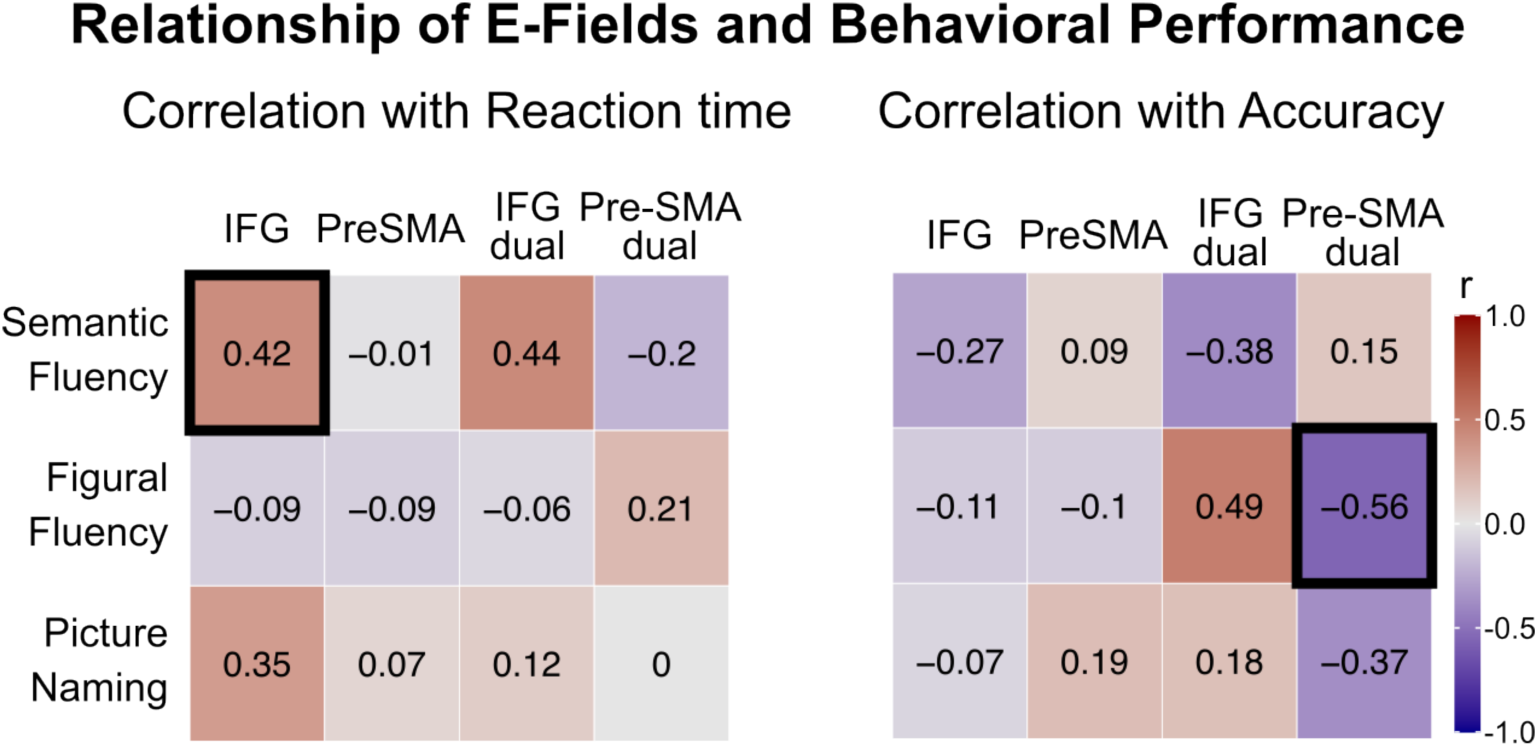
Correlation matrices for all tasks and stimulation sessions for reaction time and accuracy, respectively. Significant results after FDR correction are marked in bold. Note that for the dual-site stimulation, results are based on a regression model using both e-fields as predictors.

**Table S3.** Comparison of e-field values in subregions of cortical stimulation targets.

| Predictors | E-Field 95th Percentile |  |  |
| --- | --- | --- | --- |
|  | Estimates (CI) | Statistic | p |
| ROI [BA45 vs. BA44] | 5.65<br>(2.78 – 8.53) | 3.88 | **<0.001** |
| ROI [left pre-SMA vs. BA44] | -13.52<br>(-16.40 – -10.65) | -9.28 | **<0.001** |
| ROI [right pre-SMA vs. BA44] | -16.87<br>(-19.74 – -13.99) | -11.57 | **<0.001** |
| Session [Pre-SMA vs. IFG] | -17.47<br>(-19.50 – -15.43) | -16.95 | **<0.001** |
| ROI [BA45] x Session [Pre-SMA] | -6.43<br>(-12.18 – -0.67) | -2.20 | **0.029** |
| ROI [left pre-SMA] x Session [Pre-SMA] | 82.43<br>(76.68 – 88.19) | 28.27 | **<0.001** |
| ROI [right Pre-SMA] x Session [Pre-SMA] | 78.19<br>(72.43 – 83.94) | 26.82 | **<0.001** |
| Marginal R<sup>2</sup> / Conditional R<sup>2</sup> | 0.855 / 0.928 |  |  |

Note. All reported p values are FDR-corrected. Significant effects are marked in bold. CI: Confidence interval.

**Figure S4.**
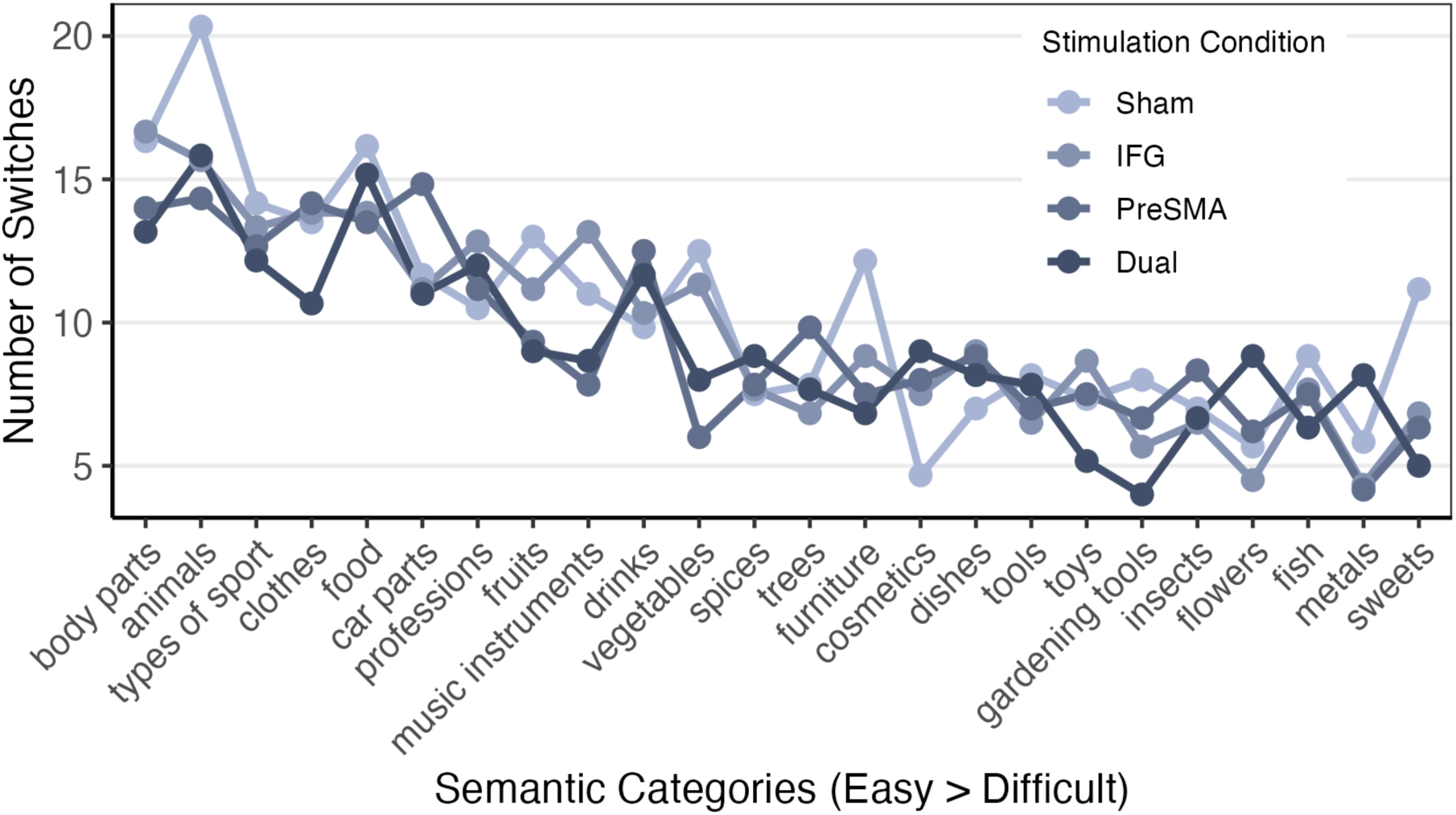
Number of switches for each semantic category and stimulation condition, ordered according to increasing difficulty.

**Table S4.** Results from mixed-effects regression for number of switches during semantic fluency.

| Number of switches per category |  |  |  |
| --- | --- | --- | --- |
| Predictors | Incidence Rate Ratios (CI) | Statistic | p |
| Target [IFG vs. Sham] | 0.93 (0.86 – 1.01) | -1.73 | 0.143 |
| Target [PreSMA vs. Sham] | 0.90 (0.83 – 0.98) | -2.44 | **0.039** |
| Target [Dual site vs. Sham] | 0.87 (0.81 – 0.95) | -3.24 | **0.007** |
| Session [2 vs. 1] | 1.06 (0.98 – 1.15) | 1.40 | 0.163 |
| Session [3 vs. 1] | 1.06 (0.97 – 1.15) | 1.28 | 0.200 |
| Session [4 vs. 1] | 1.08 (1.00 – 1.18) | 1.94 | 0.053 |
| Category type [Hard vs. Easy] | 0.60 (0.52 – 0.69) | -7.28 | **<0.001** |
| Marginal R<sup>2</sup> / Conditional R<sup>2</sup> | 0.255 / 0.572 |  |  |

Note. Reported p-values of stimulation conditions are FDR-corrected. Significant effects are marked in bold. CI: Confidence interval.

